# Shear stress targeted delivery of nitroglycerin to brain collaterals improves ischaemic stroke outcome

**DOI:** 10.1101/2025.03.25.644833

**Authors:** Magdalena Litman, Sara Azarpeykan, Rebecca J Hood, Kristy Martin, Debbie Pepperall, Daniel Omileke, Oktay Uzun, Deen Bhatta, Yuen K Yong, Alex Chan, Nicholas Hough, Sarah Johnson, Pablo Garcia Bermejo, Ferdinand Miteff, Carlos Garcia Esperon, Yvonne Couch, Alastair M Buchan, Neil J Spratt, Netanel Korin, Donald E Ingber, Daniel J Beard

**Affiliations:** School of Biomedical Science and Pharmacy, The University of Newcastle, Newcastle, Australia; Discipline of Anatomy and Pathology, School of Biomedicine, Faculty of Health and Medical Sciences, The University of Adelaide, Australia; Heart and Stroke Programme, Hunter Medical Research Institute, Newcastle, Australia; Wyss Institute of Biologically Inspired Engineering, Harvard University, Boston, MA, USA; School of Engineering, University of Newcastle, Callaghan, NSW 2308, Australia; Department of Neurology, John Hunter Hospital, University of Newcastle, Australia; Acute Stroke Programme, Radcliffe Department of Medicine, University of Oxford, Oxford, UK; Department of Biomedical Engineering, Technion-Israel Institute of Technology, Haifa, Israel; Vascular Biology Program and Department of Surgery, Children’s Hospital Boston and Harvard Medical School, Boston, MA, USA; Harvard John A. Paulson School of Engineering and Applied Sciences, Cambridge, MA, USA

## Abstract

In patients with ischaemic stroke, retrograde perfusion of the penumbra by the leptomeningeal collateral vessels (LMCs) is a strong predictor of clinical outcome, thus raising the possibility that enhancing LMC flow could offer a novel therapeutic approach. Here, using computational modelling we show that LMCs experience elevated fluid shear stress that is significantly higher than that in other blood vessels during ischaemic stroke in animals and humans. We take advantage of this to selectively enhance flow in LMCs using shear-activated nanoparticle aggregates carrying the vasodilator nitroglycerin (NG-NPAs) that specifically release drug in regions of vessels with high shear stress (≥100 dyne/cm^2^). The NG-NPAs significantly increased LMC-mediated penumbral perfusion, decreased infarct volume, and reduced neurological deficit without altering systemic blood pressure in a rat ischaemic stroke model. The NG-NPAs also did not cause known common side effects of systemic nitrate administration, such as systemic hypotension, cerebral vascular steal, cortical vein dilation, or intracranial pressure elevation. Systemic administration of free NG at the maximal tolerated dose, which was ten times higher than the dose of NG used in the NG-NPAs, did not enhance LMC perfusion and dropped blood pressure. Thus, packaging NG within shear-activated NPAs can potentially enable this widely available vasodilator to become a highly effective therapeutic for ischaemic stroke.

## Introduction

Stroke, resulting in disruption of blood flow to the brain, is the 3^rd^ leading cause of death and a major cause of long-term disability worldwide^1^. Ischaemic stroke, which accounts for 71% of all strokes globally, is caused by cerebral arterial occlusion^2^. The goal of acute ischaemic stroke therapies is to salvage the ischaemic penumbra, which is the region of brain tissue beyond the occlusion that is functionally compromised but remains potentially salvageable. The penumbra receives retrograde blood flow via cerebral collateral vessels^3^ in the brain’s leptomeningeal collateral (LMC) circulation that are formed through end-to-end anastomoses of distal branches of adjacent arterial territories. The LMCs provide residual retrograde cerebral blood flow when conventional routes of arterial supply are compromised^4^.

Importantly, advanced clinical imaging over the past decade has led to recognition of the key importance of LMCs in ischaemic stroke outcome^5^. The presence of good LMC perfusion is associated with a larger penumbra, better rates of reperfusion, and improved functional outcomes following both thrombolysis and thrombectomy^6–8^. This is most likely because patients with better collaterals have a large volume of penumbra when they arrive at hospital^9,10^. LMCs have such a large influence on response to reperfusion therapies and stroke outcome, that LMC status and penumbral volume are now used to select patients most likely to benefit from reperfusion therapies^11,12^. However, a large portion of stroke patients have co-morbidities such as hypertension (in ∼55% of stroke patients), which is associated with poor LMC status. These patients may therefore benefit greatly from therapeutic interventions that enhance LMC blood flow^13^. While the potential benefits of therapeutic approaches that enhance collateral flow and thereby improve stroke outcome are widely recognised, finding an effective method to enhance collateral flow has proved to be very challenging^14^.

One of the greatest obstacles to developing collateral therapies is the inability to selectively vasodilate LMCs without causing systemic vasodilation and hypotension^15^. Healthy cerebral vasculature can autoregulate (vasodilate) in response to low blood pressure and maintain normal perfusion^16^. However, collaterals in the ischaemic penumbra have already reached their limits of autoregulation, and so blood flow becomes entirely dependent on cerebral perfusion pressure^17^. Thus, systemic vasodilators that lower blood pressure may have a “reverse Robin Hood” effect - redirecting or stealing blood flow away from the ischaemic brain territory^18^. Further, systemic vasodilators have a higher affinity for cerebral veins, leading to venodilation and increased intracranial pressure, which further reduces cerebral perfusion pressure^15^. As a result, clinical trials of systemic vasodilators in stroke showed no change in penumbral blood flow^19^ and no improvement in patient outcome^20^. Based on this evidence, the American Heart Association Guidelines advise against use of systemic vasodilatory agents for ischaemic stroke treatment^21^.

This present study is based on the hypothesis that selective dilation of LMCs may be achieved by targeting vasodilators to these vessels based on work which has revealed that LMCs have unique blood flow haemodynamics that are potentially exploitable therapeutically^22^. More specifically, when the middle cerebral artery is occluded in rats, the large pressure differential between the occluded and patent vascular territories causes a 10-fold increase in blood flow velocity through LMCs and an associated increase in fluid shear stress (>100 dyne/cm^2^) selectively within these vessels^23,24^. Importantly, this level of shear stress is significantly higher than that in other vascular beds in the body^25,26^. Shear stress may aid in the vasodilation of LMCs to promote perfusion to the penumbra^27^; however, hypertensive animals exhibit both poorer baseline LMC blood flow and impaired flow-mediated vasodilation^24,28^, ultimately leading to poor penumbral perfusion. These observations raise the possibility that if one could selectively target a vasodilator to LMCs that experience high shear stress, this could provide a way to enhance LMC flow and thereby, reduce stroke morbidity and mortality.

To approach this challenge, we leveraged a previously described drug delivery platform composed of spray-dried, drug-loadable, polymeric nanoparticles (∼180 nm diameter) that are formed into platelet sized (∼3 μm diameter) nanoparticle aggregates (NPAs). Importantly, these NPAs selectively release the nanoparticles and their contents where they experience pathological fluid shear stress (>100 dyne/cm^2^) when injected systemically into the circulation^29^. When these shear-activated NPAs were loaded with tissue plasminogen activator (tPA) and injected intravenously, they dissolved pulmonary emboli using one hundredth the dose of soluble tPA required to accomplish the same response in a mouse model^29^. These tPA-loaded NPAs also enhanced removal of blood clots in large brain arteries through enzymatic dissolution^30^. In the present study, we loaded these shear-activated NPAs with nitroglycerin (NG-NPAs) to explore whether we could selectively target this potent vasodilator to the smaller LMCs surrounding the penumbra in the context of ischaemic stroke. Our results show that NG-NPAs can significantly enhance LMC blood flow, improve perfusion of the penumbra during stroke, decrease infarct volume, and reduce neurological deficit without causing systemic side effects that limit the use of free nitroglycerin for this life-threatening condition.

## Results

### Increased shear stress in LMCs in rats and humans

To determine the translational potential of our shear targeted drug delivery concept, we quantified fluid shear stress levels in collateral vessels of both healthy normotensive Wistar rats and spontaneously hypertensive rats (SHRs). We included hypertensive rats because high blood pressure is one of the most common co-morbidities and leading modifiable risk factor for stroke, and it is associated with LMC blood flow^31,32^. We found that before the stroke, LMC blood flow was bi- directional and very slow. However, upon induction of stroke by blocking the proximal segment of the middle cerebral artery (MCA), flow became unidirectional.

The blood flowed from the anterior cerebral artery (ACA) towards the MCA and the velocity increased dramatically while vessel diameter only increased slightly. When we quantified shear stress in LMC following experimental stroke, we detected significant (5- to 6-fold) increases in both normotensive Wistar rats (**Fig. 1a**) and SHRs (**Fig. 1b**).

**Fig. 1.**
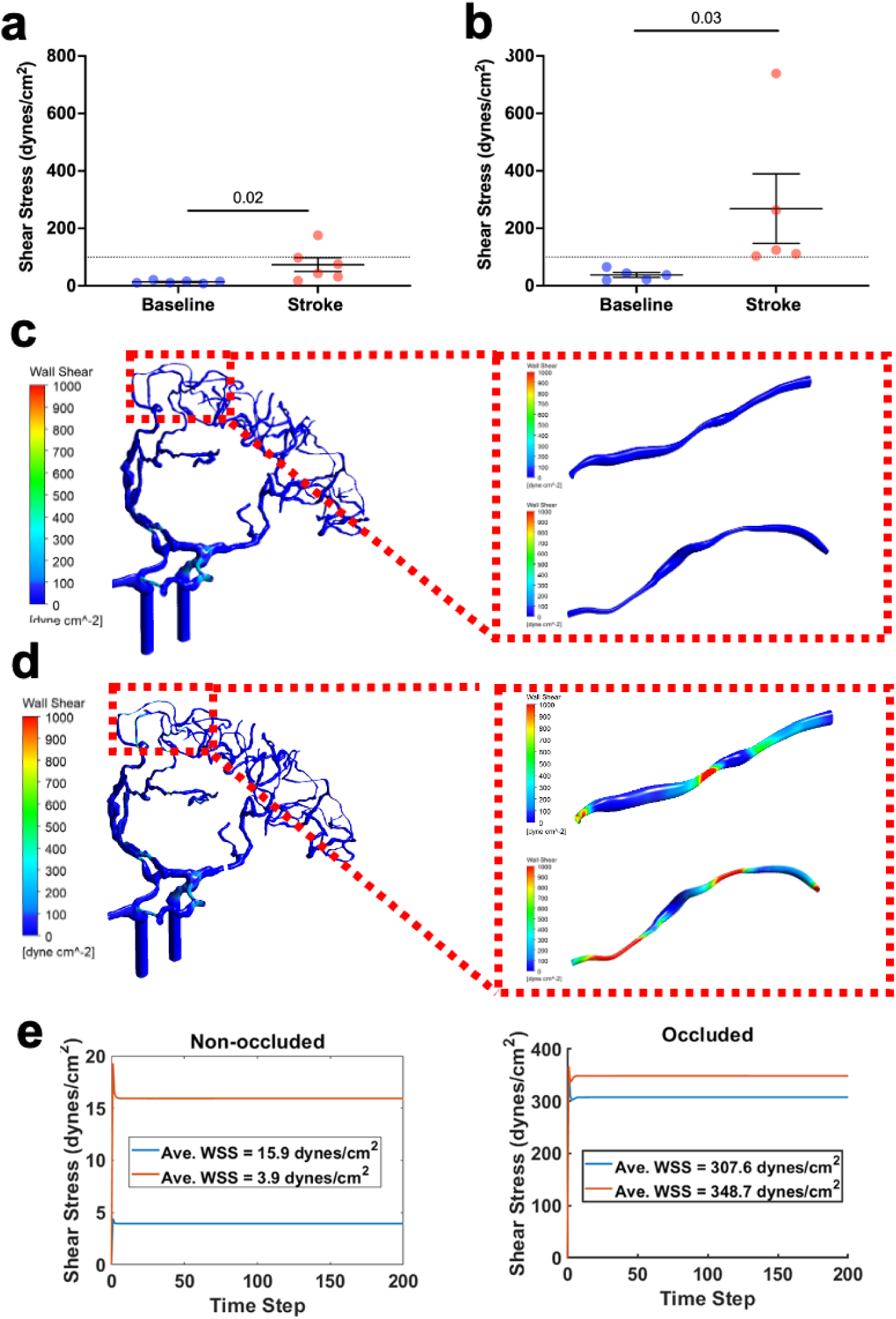
Shear stress is significantly increased in LMC during stroke. Average shear stress in collateral vessels in Wistar rats **(a)** and spontaneous hypertensive rats **(b)** at baseline versus stroke. **(c)** Computational Fluid Dynamic (CFD) modelling of wall shear stress in human cerebral vessels without proximal occlusion of the MCA (unoccluded model), with more detailed wall shear stress analysis in individual collateral vessels (inset). **(d)** Similar CFD modelling proximal occlusion of the MCA (occluded stroke model). **(e)** Quantified average wall shear stress over time in individual collateral vessels (the wall shear stress in each collateral is represented by a red and blue line) from the unoccluded versus occluded model from **c** and **d**, respectively. The wall shear stress is colour coded to represent dyne/cm^2^ as per the scale. One-sided, Wilcoxon test was used to compare average shear before and after stroke in animal studies. Presented values represent mean ± SEM.

### Computational fluid dynamic analysis of blood flow in human LMCs

To explore the potential clinical relevance of this observation, we carried out computational fluid dynamics simulations of the LMCs following a simulated ischaemic stroke in a human. CT angiogram images of a patient who had an ischaemic stroke with proximal MCA occlusion were segmented to produce a 3D model of the ACA, MCA, and LMCs that link them before (**Fig. 1c)** and after **(Fig. 1d**) occlusion. When we simulated shear stress in two LMCs (the shear stress in each collateral is represented by a red and blue line in **Fig.1e**) under non-occluded and occluded (ischaemic stroke following MCA occlusion) conditions, we found that there were 20- to 60-fold increases in shear stress following occlusion of the LMCs (**Fig. 1e**).

### Shear-activated delivery of nitroglycerin selectively increases LMC perfusion

As hypertensive patients are the most difficult to treat and we observed the greatest level of shear stress in SHRs, all further experiments were conducted using the hypertensive animals. MCA occlusion was induced for 70 min in male SHRs and laser speckle contrast imaging was used to measure tissue perfusion in the LMCs (**Fig. 2a**). Animals were randomised to receive intravenous infusion of NPAs without drug (Blank-NPAs) or NG-NPAs (4 μg NG/kg/min) commencing 25 minutes after the MCA was blocked, and infarct volume was measured at 24 h (**Fig. 2a**). These studies revealed that infusion of NG-NPAs increased LMC perfusion by almost 45% between 12 and 40 min compared to pre-infusion baseline, while there was no significant change when the same dose of Blank-NPAs were administered (**Fig. 2b**). The increase in LMC perfusion averaged over the entire infusion period (**Fig. 2c**) and the peak level of perfusion (**Fig. 2d**) for the NG-NPAs were also significantly higher than that of the Blank-NPAs, which again did not produce a significant change compared to baseline.

**Fig. 2.**
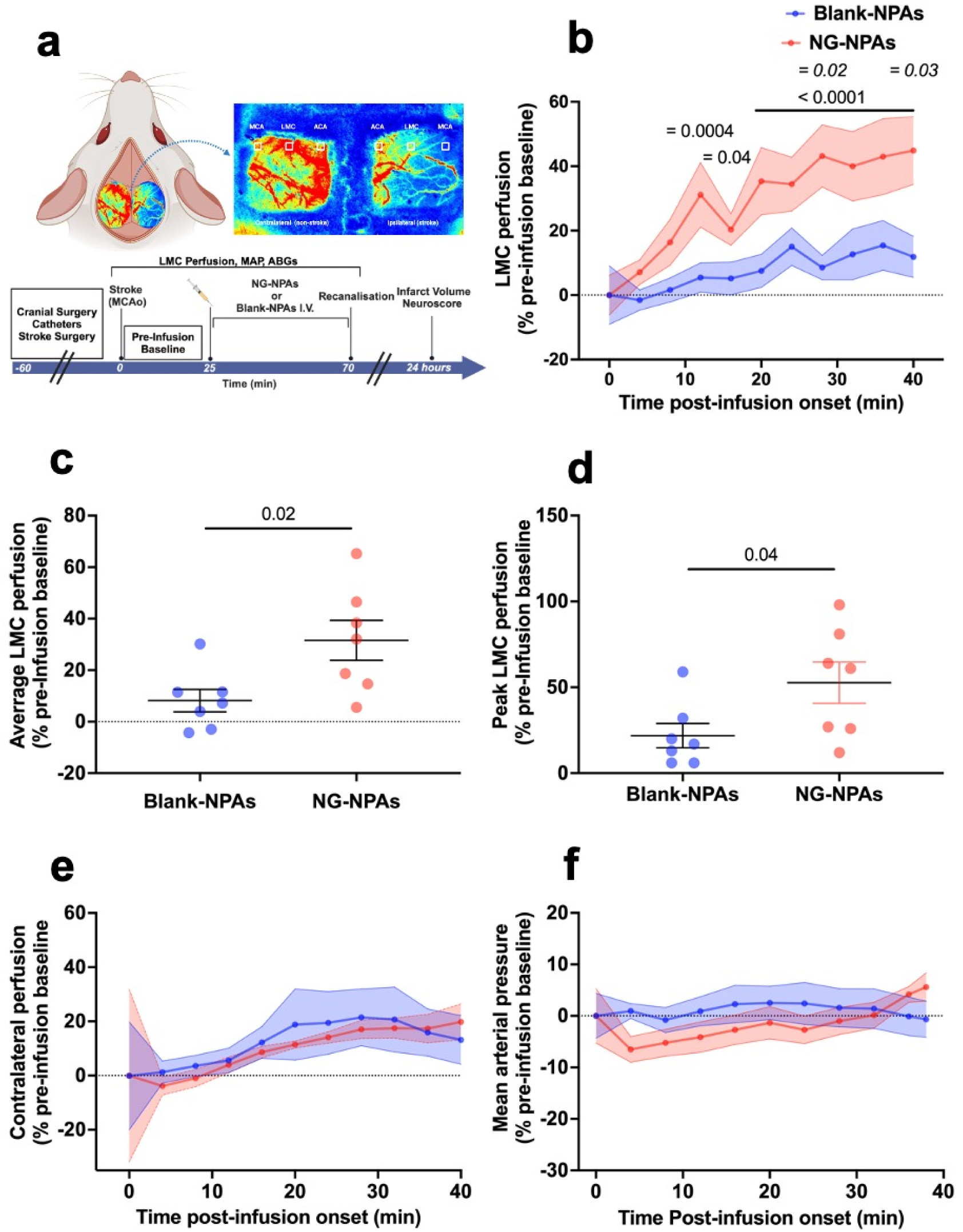
NG-NPA infusion significantly increases LMC perfusion without causing changes in contralateral collateral perfusion or a drop in mean arterial pressure in spontaneously hypertensive rats. **(a)** Diagram demonstrating perfusion regions analysed using laser speckle contrast imaging during experimental stroke and experimental timeline. **(b)** LMC perfusion in the stroke region in NG-NPA (red) treated vs. Blank-NPA (blue) treated SHRs, calculated as a % change from pre-drug infusion baseline. We conducted a repeated measure 2-Way ANOVA to assess the effect of NG-NPA vs. Blank-NPA treatment over time. F (1, 12 ) = 6.936, p = 0.02 for treatment, F (10, 120) = 8.035, p < 0.0001 for time, F (10, 120) = 2.1, p= 0.03 for interaction. Sidak’s post-test was used to compare NG-NPA vs. Blank-NPAs at different time-points (p values on graph in italics, p < 0.05 at 30 and 40 min). Dunnett’s post-test was used to compare each time-point back to pre-infusion baseline in NG-NPA and Blank NPA groups (p values on graph in normal text, NG-NPA: p < 0.05 at 16 min; p < 0.001 at 12 min; p < 0.0001 between 20 and 40 min). **(c)** Average LMC perfusion in the stroke region. We used an un-paired t-test to assess differences between NG-NPAs (red dots) vs. Blank-NPAs (blue dots). T (12) = 2.63, p = 0.02 vs. Blank-NPAs. **(d)** Peak LMC perfusion in the stroke region. We used an un-paired t-test to differences between NG-NPAs (red dots) vs. Blank-NPAs (blue dots). T (12) = 2.212, p = 0.04 vs. Blank-NPAs. **(e)** LMC perfusion in the contralateral (control) hemisphere in NG-NPA (red) treated vs. Blank-NPA (blue) treated SHRs. We conducted a repeated measure 2-Way ANOVA to assess the effect of NG-NPA vs. Blank-NPA treatment over time. F (1,12 ) = 0.156 , p = 0.69 for treatment, F (10, 120) = 1.35 , p = 0.21 for time, F (10, 120) = 0.08 , p > 0.99 for interaction. **(f)** Mean Arterial Pressure in NG-NPA (red) treated vs. Blank-NPA (blue) treated. We conducted a repeated measure 2-Way ANOVA to assess the effect of NG-NPA vs. Blank-NPA treatment over time. F (1,12 ) = 0.49 , p = 0.5 for treatment, F (10, 120) = 1.02 , p = 0.42 for time, F (10, 120) = 1.49 , p = 0.15 for interaction. Values represent mean ± SEM.

In contrast, systemic administration of the shear targeted NG-NPAs did not significantly alter perfusion in the LMC region in the contralateral (non-stroke/control) hemisphere compared to pre-infusion baseline or relative to Blank-NPAs (**Fig. 2e**).

Importantly, infusion of the NG-NPAs also did not reduce systemic blood pressure relative to pre-infusion baseline or Blank-NPAs (**Fig. 2f**). All other physiological variables were within physiological range and were not significantly different between treatment groups (**Table 1**). Thus, the shear-activated NG-NPAs appeared to selectively produce vasodilation and enhanced perfusion precisely where our rat experiments and human models predicted fluid shear stress to be preferentially elevated.

**Table 1.** Physiological parameters measured pre-and-during NG-NPA vs Blank-NPA infusion from Study 1. Mean ± SEM of MAP, mean arterial pressure; RR, respiratory rate; HR, heart rate; SPO_2,_ oxygen saturation; paO_2,_ arterial partial pressure of oxygen; paCO_2_, arterial partial pressure of carbon dioxide.

|  | NG-NPAs |  | Blank-NPAs |  |
| --- | --- | --- | --- | --- |
|  | Pre-infusion | During infusion | Pre-infusion | During infusion |
| MAP (mmHg) | 158.7 ± 10.56 | 155 ± 10.27 | 158.8 ± 8.18 | 171.7 ± 7.36 |
| RR (BMP) | 52.14 ± 2.77 | 53.10 ± 2.68 | 45.21 ± 2.09 | 44.88 ± 2.87 |
| HR (BPM) | 398.4 ± 12.28 | 407.3 ± 14.86 | 393.6 ± 11.91 | 417.3 ± 12.86 |
| SPO2 (%) | 95.89 ± 1.03 | 97.11 ± 1.06 | 95.74 ± 0.72 | 96.5 ± 0.54 |
| paO2 (mmHg) | 97.25 ± 8.11 | 99.50 ± 5.87 | 114.5 ± 6.84 | 112.5 ± 9.18 |
| paCO2 (mmHg) | 43.40 ± 0.26 | 41.45 ± 0.71 | 41.03 ± 1.53 | 39.65 ± 1.14 |
| pH | 7.387 ± 0.0056 | 7.399 ± 0.007 | 7.414 ± 0.01 | 7.39 ± 0.009 |

### Shear-targeted delivery of nitroglycerin improves stroke outcomes

At 24 h after stroke onset, animals underwent assessment of neurological deficits, and they were then euthanised, perfusion fixed, and brains were collected for coronal sectioning and histological evaluation of infarct size using haematoxylin and eosin staining. Quantification of infarct size revealed that NG-NPAs significantly reduced final infarct volume by over 35% compared to Blank-NPAs (**Fig. 3a**). Infarct probability maps also indicated that a reduction in the frequency of infarcts in the cortex located downstream of the LMC (i.e. in the penumbral region) is likely responsible for the reduction in overall infarct volume in the NG-NPA group (**Fig. 3b**). There was a very large and significant inverse correlation between average level of LMC perfusion and infarct volume (spearman’s r = -0.69, p = 0.01; **Fig. 3c**). The neurological deficit scores also were significantly lower in animals receiving NG- NPAs (median score = 2; range 1-4) versus Blank-NPAs (median = 4; range 3-5) (**Fig. 3d**).

**Fig. 3.**
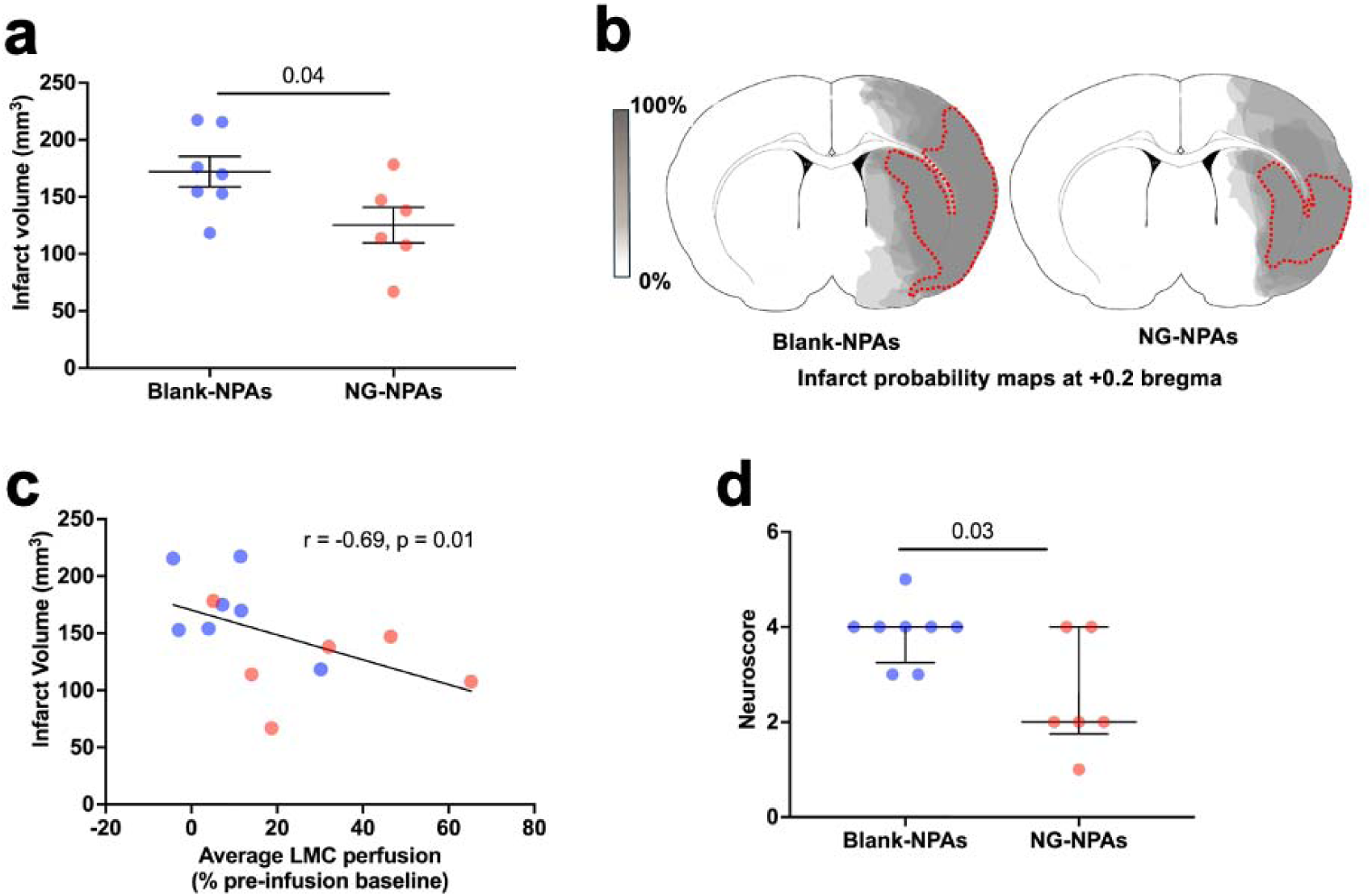
NG-NPAs significantly reduced infarct volume and improved neurological function 24 h post stroke in spontaneously hypertensive rats. **(a)** Infarct volume was assessed histologically at 24 hours after stroke onset. We used an un-paired t-test to differences between NG-NPAs (red dots) vs. Blank-NPAs (blue dots). T (11) = 2.29, p = 0.04 vs. Blank-NPAs. **(b)** Infarct probability maps were generated from infarct areas taken from the brain region in the central region supplied by the middle cerebral artery (+0.2 bregma brain slice). Darker grey regions represent areas with a higher probability of infarction. **(c)** Spearman’s correlation between infarct volume and average LMC perfusion. **(d)** Neurological score assessing forelimb flexion, lateral push and torso twisting (0 = normal, 6 = maximal deficit). We used a Mann-Whitney U test to to assess differences between NG-NPAs (red dots) vs. Blank-NPAs (blue dots). U = 9, p = 0.03. Values represent mean ± SEM.

### Shear-targeted delivery did not cause detectable adverse effects

Nitroglycerin (NG) administration as a free drug has previously been shown to have adverse effects due to dilation of cerebral veins, which increases intracranial pressure^15^. Importantly, when we infused the NG-NPAs, we did not detect any change in cortical vein diameter compared to pre-infusion baseline or relative to administration of Blank-NPAs (**Fig. 4a**). The NG-NPA treatment also did not change intracranial pressure compared to pre-infusion baseline (**Fig. 4b**) and it was significantly lower in the NG-NPA group compared to Blank-NPAs at all time-points, including pre-infusion baseline (**Fig. 4b**).

**Fig. 4.**
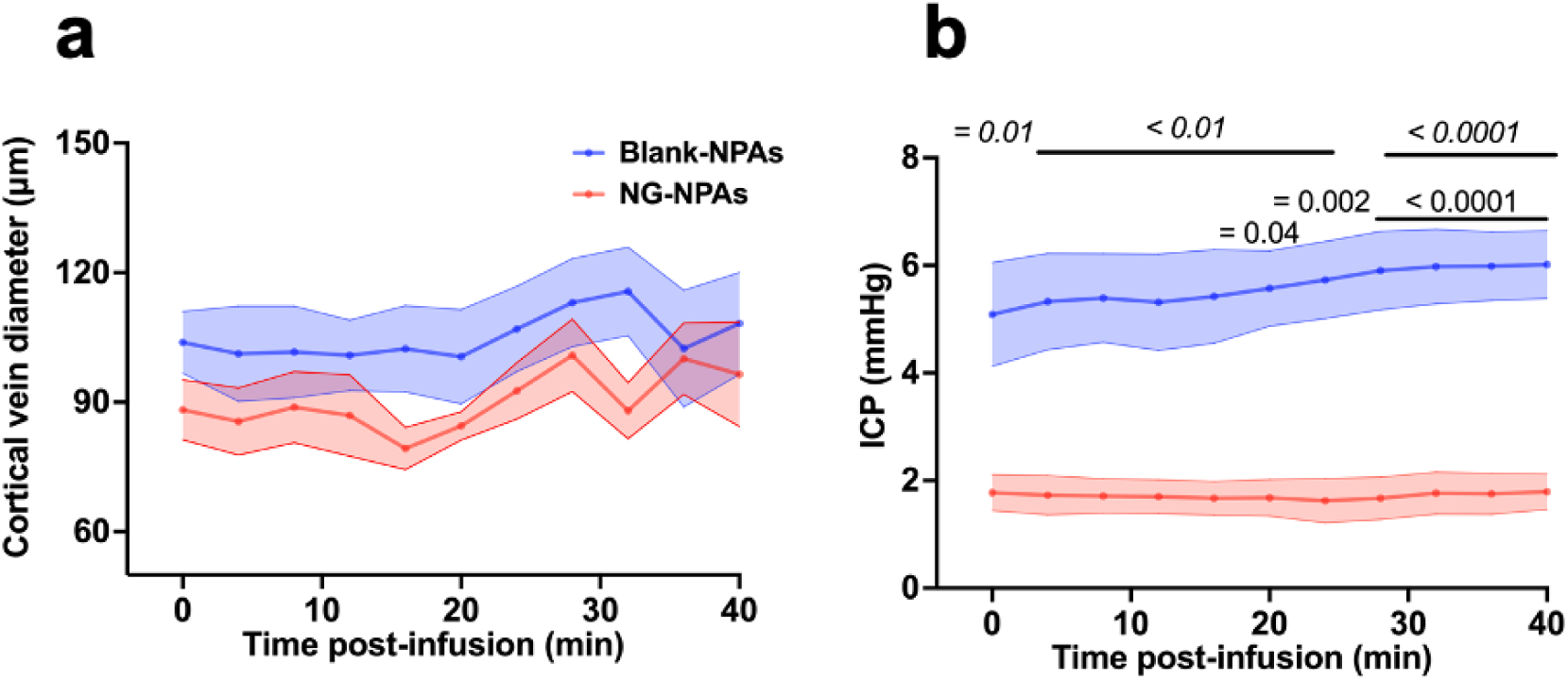
NG-NPAs did not cause changes in cortical vein diameter or intracranial pressure in spontaneously hypertensive rats. **(a)** Cerebral vein diameter following NG-NPA (red dots) or Blank-NPA (blue dots) administration. We conducted a Mixed-Effects Model to assess the effect of NG-NPA vs. Blank-NPA treatment over time. p = 0.2 for treatment, p = 0.002 for time, p= 0.4 for interaction. **(b)** Intracranial pressure following NG-NPA (red dots) or Blank-NPA (blue dots) administration. We conducted a repeated measure 2-Way ANOVA to assess the effect of NG-NPA vs. Blank-NPA over time. F (1, 5 ) = 16.38, p = 0.001 for treatment, F (10, 50) = 3.7, p < 0.0009 for time, F (10, 50) = 3.428, p= 0.002 for interaction. Sidak’s post-test was used to compare NG-NPA vs. Blank-NPAs at different time-points (p values on graph in italics, p = 0.01 at 0 min; p < 0.01 between 4 and 24 min and p < 0.0001 between 28 and 40 min). Dunnett’s post-test was used to compare each time-point back to pre-infusion baseline in NG-NPA and Blank NPA groups (p values on graph in normal text, Blank-NPAs: p = 0.04 at 20-min; p = 0.002 at 24 min and p < 0.0001 between 28 and 40 min). Values represent mean ± SEM.

### Administration of free NG is ineffective

We next wanted to confirm that the LMC perfusion-increasing effects of the shear-targeted NG-NPAs we observed could not be obtained using a similar dose of free NG (4 μg/kg/min). Infusion of the same dose of free NG did not significantly change LMC perfusion relative to pre-infusion baseline or to saline controls in the region of the stroke (**Fig. 5a**) or in the LMCs within the contralateral (non-stroke/control) hemisphere (**Fig. 5c**). Even when we administered 10 times the dose of free NG (40 μg/kg/min), we could not detect a significant change in perfusion in LMCs in the stroke area (**Fig. 5b**) or in the LMCs in the contralateral hemisphere (**Fig. 5d**).

**Fig. 5.**
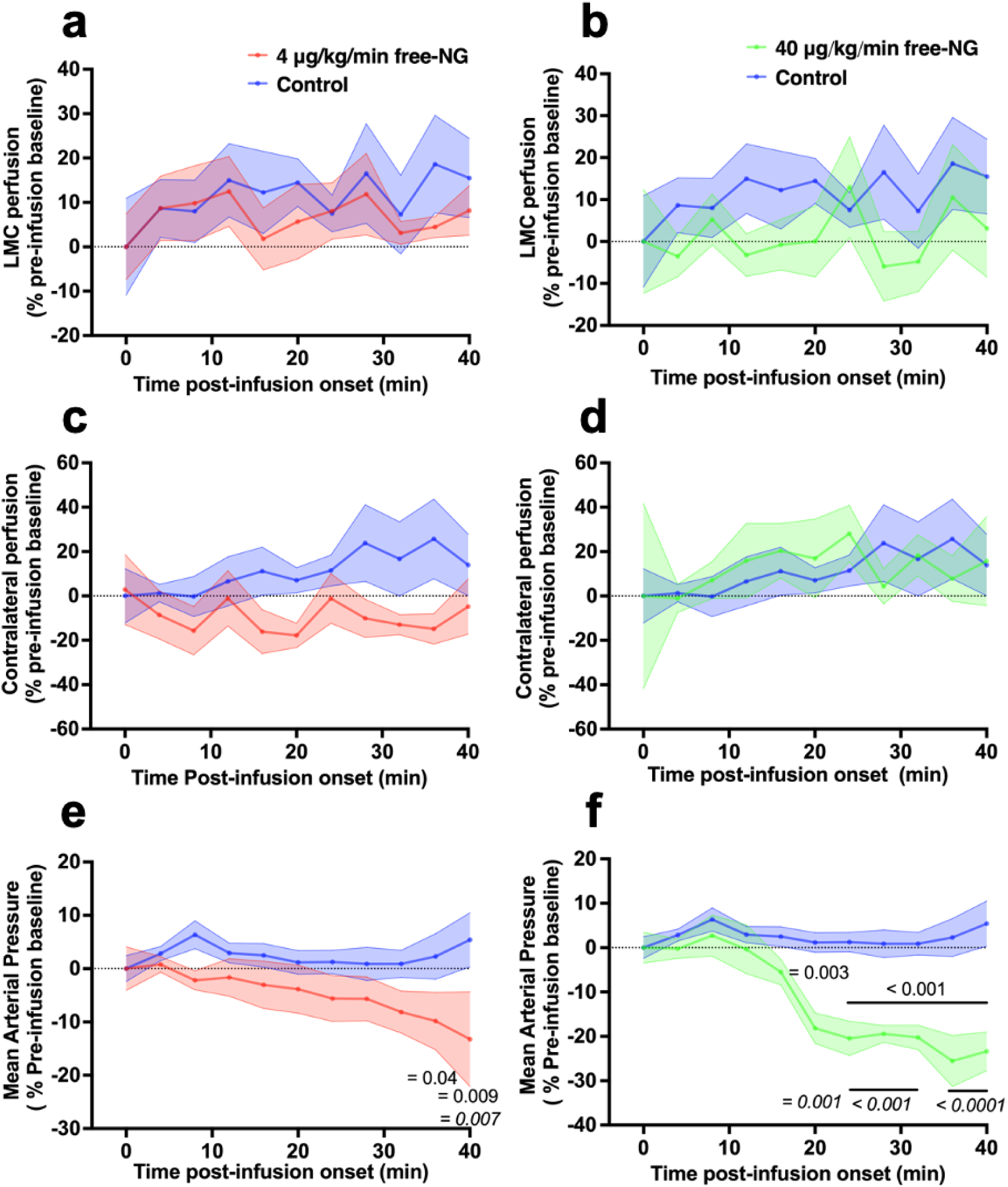
Administration of free-NG did not improve LMC perfusion but caused a drop in mean arterial pressure. **(a)** The effect of 4μg/kg/min free-NG on ipsilateral LMC perfusion. We conducted a Mixed-Effects Model to assess the effect of 4μg/kg/min free-NG vs. saline control over time. p = 0.69 for treatment, p = 0.52 for time, p= 0.98 for interaction. **(b)** The effect of a 10 times higher dose (40μg/kg/min) of free-NG on ipsilateral LMC perfusion. We conducted a Mixed-Effects Model to assess the effect of 40μg/kg/min free-NG vs. saline control over time. p = 0.22 for treatment, p = 0.8 for time, p = 0.85 for interaction. **(c)** The effect of 4μg/kg/min free-NG on contralateral perfusion. We conducted a Mixed-Effects Model to assess the effect of 4μg/kg/min free-NG vs. saline control over time. p = 0.09 for treatment, p = 0.96 for time, p= 0.66 for interaction. **(d)** The effect of high dose (40μg/kg/min) free-NG on contralateral perfusion. We conducted a Mixed-Effects Model to assess the effect of 40μg/kg/min free-NG vs. saline control over time. p = 0.82 for treatment, p = 0.83 for time, p= 0.96 for interaction. **(e)** The effect of 4μg/kg/min free-NG on mean arterial pressure. We conducted a Mixed-Effects Model to assess the effect of 4μg/kg/min free-NG vs. saline control over time. p = 0.07 for treatment, p = 0.89 for time, p = 0.04 for interaction. Sidak’s post-test was used to compare 4μg/kg/min free-NG vs. saline control at different time-points (p values on graph in italics, p = 0.007 at 40 min). Dunnett’s post-test was used to compare each time-point back to pre-infusion baseline in 4μg/kg/min free-NG and saline control groups (p values on graph in normal text, 4μg/kg/min free-NG: p = 0.04 at 36 min; p = 0.009 at 40 min). **(f)** The effect of 40μg/kg/min free-NG on mean arterial pressure. We conducted a repeated measure 2-Way ANOVA to assess the effect of 40μg/kg/min free-NG vs. saline control over time. p = 0.002 for treatment, p < 0.0001 for time, p < 0.0001 for interaction. Sidak’s post-test was used to compare 40μg/kg/min free-NG vs. saline control at different time-points (p values on graph in italics, p = 0.001 at 20 min; p < 0.001 between 24 and 32 min; p < 0.0001 between 36 and 40 min). Dunnett’s post-test was used to compare each time-point back to pre-infusion baseline in 40μg/kg/min free-NG and saline control groups (p values on graph in normal text, 40μg/kg/min free-NG: p = 0.0003 at 20 min; p < 0.0001 between 24 and 40 min). Values represent mean ± SEM.

Most concerning was that infusion of an equivalent amount of free NG significantly lowered systemic blood pressure 40 min post-infusion relative to pre-infusion baseline (**Fig. 5e**), as observed in patients who received systemic administration of NG for treatment of stroke^20^. As expected, this was further exacerbated by administration of the 10-fold higher dose of free NG (**Fig. 5f**). Thus, this animal model replicates clinical observations seen in stroke patients treated with free NG, and it shows that free NG cannot provide therapeutic effects for ischaemic stroke like those produced when NG is targeted to regions of high shear stress using the NPA delivery system.

## DISCUSSION

Modulation of LMC perfusion has been recognized as a potentially exciting therapeutic approach to ischaemic stroke in human patients; however, producing this response without dangerous side effects has proven elusive. A key reason has been the inability to selectively enhance perfusion within relevant LMCs while avoiding systemic vasodilation and resultant hypotension. In this study, we showed that fluid shear stress in LMC vessels in the region downstream of the occlusion is significantly higher during ischaemic stroke in both normotensive and hypertensive animals, as well as in humans based on computational modelling. Moreover, we show that this hemodynamic peculiarity can be leveraged using a shear-activated nanoparticle drug delivery system to selectively deliver a potent FDA-approved vasodilator (NG) to these sites and thereby, enhance penumbral perfusion, while avoiding deleterious side effects.

This mechanotherapeutic approach is supported by the finding that fluid shear stress increases abruptly immediately after the MCA is experimentally occluded in normotensive and hypertensive rats as shown here, as well as in normotensive mice^33^. However, fluid shear stress measured in the LMCs of the hypertensive rats was much higher than that in normotensive animals, increasing to values well above the threshold of activation for NG-NPAs to release their contents (>∼100 dyne/cm^2^). This is highly relevant from a translational perspective, as the normotensive Wistar rats we used, have larger diameter LMC vessels and smaller infarcts than the SHRs.^34,35^ Hypertension is also seen in the majority of stroke patients^36^, and it is associated with poor collateral blood flow and worse outcomes in both patients^37^ and animals^38^. Therefore, hypertensive patients would be more likely to respond to and have the most to gain from the NG-NPA treatment, therefore this could be a choice patient population for future clinical studies.

As a control in this study, we measured perfusion through LMCs in the same animal on the contralateral side of the brain that did not experience stroke and found that infusion of the NG-NPAs had no effect on their perfusion. Most importantly, the specific enhancement of perfusion through the stroke associated LMCs resulted in a significant reduction in infarct size, consistent with clinical observations suggesting that better collateral perfusion is associated with smaller infarct volumes^39^. Our findings also demonstrated that targeted delivery of NG to the LMCs in stroke regions using the shear-activated NPA delivery platform led to selective collateral vasodilation without producing systemic hypotension or increased intracranial pressure. These results suggest that systemic arterial vasodilation and cerebral venodilation also did not occur.

Systemically administered NG has been previously proposed as a potential therapeutic agent in the treatment of acute ischaemic stroke and it has been administered as a transdermal patch in seven randomised clinical trials^40–46^. The recently published RIGHT-2 trial (2019) showed that transdermal NG was effective in lowering systemic blood pressure but was neutral for functional outcome at 3 months post-stroke. In fact, the point estimates indicated a trend for harm in patients with the final diagnosis of stroke or transient ischaemic attack^43^. As perfusion imaging was not performed on trial patients, the reasons for these results are unclear. A possible explanation for this is that the drop in systemic blood pressure counteracted the benefits of cerebral vessel dilation caused by NG. This is supported by the findings from another study measuring changes in cerebral perfusion associated with transdermal NG treatment. Of the 18 patients treated with transdermal NG within 5 days of symptoms onset, blood pressure was reduced, and cerebral blood flow remained unaffected^40^. A subsequent perfusion study using the same treatment protocol as the RIGHT-2 trial showed no improvement in penumbral perfusion on imaging, suggesting that the drug might not have made it to the brain or, if it did, the therapeutic concentration in the brain was not reached, resulting in no change in cerebral perfusion^47^. Thus, our finding that intravenous infusion of free NG using an equivalent dose to what was in the NPAs did not enhance collateral perfusion, but caused systemic vasodilation and systemic hypotension, is consistent with known effects observed in human patients^45,48^. Furthermore, administration of a 10 times higher dose of free NG did not affect tissue perfusion in this experimental model, but significantly lowered systemic blood pressure, suggesting that any effect of dilation of collaterals on blood flow was counteracted by the reduction in blood pressure. Because of these dose-limiting hypotensive effects, we were not able to find a dose of free NG that produced the equivalent collateral enhancing effect to NG-NPAs.

In contrast, our results suggest that by selectively targeting short-acting vasodilators to LMCs using the shear-activated NPA delivery platform, only a low dose of active NG drug is needed to increase penumbral perfusion, reduce infarct size, and decrease neurological deficit, while avoiding systemic complications. It is important to note that the total dose of NG (0.16 mg/kg) we loaded in the NG-NPAs, was much lower than that used in other preclinical studies that were looking at alternative ways to use free NG for the treatment of stroke. For example, in one study that was designed to target NG to the cerebral vasculature and limit systemic side effects, by direct intraarterial administration through the internal carotid artery following reperfusion in a mouse stroke model, they found that only the two highest doses (2 and 4 mg/kg) that were more than 10 times that dose loaded in our NG- NPAs were found to reduce infarct volume and improved functional outcome; however, even then they were not able to demonstrate improvement in cerebral blood flow^49^. Reduced infarct volume also has been shown with NG administered at a very high dose (10 mg/kg, i.p.) 20 min pre-stroke in rats, but the same study failed to show benefits with post-stroke administration^50^. Unfortunately, while overall the results were positive, the treatment window, route of administration, and very high doses of drug required in both studies are not relevant for use in clinical practice. A more recent study using treatment with a NG patch loaded with doses equivalent to human trials (0.06 mg/kg) failed to show significant improvement of cerebral blood flow, infarct size or functional outcome in a mouse model of ischaemic stroke, in line with clinical data^51^. In that study, increasing the NG doses to up to 48 mg/kg (700 times the clinical dose) did not produce any additional benefit on cerebral blood flow, infarct size or functional outcome. Furthermore, a recent systematic review and meta-analysis of these experimental studies of NG in stroke, revealed an overall neutral effect of systemic NG on infarct volume^52^. Thus, our results with NG-NPAs are both highly novel and clinically relevant.

Although our results demonstrate the ability of NG-NPAs to selectively enhance collateral-mediated penumbral perfusion in animals with ischaemic stroke and hypertension, we acknowledge that further work will be needed to confirm benefits in a more easily achievable clinical time-window. NG-NPAs were infused at 25 min post vascular occlusion because of constraints due to very rapid infarct expansion seen in SHRs. Hence, to conduct studies in the most humane manner, we did not wish to delay drug infusion to the point where it resulted in very large infarcts. However, it should be noted that this is a therapy with strong potential to be delivered in-ambulance, in which case a 25 min time interval may be achievable in some patients, and a median treatment time of 45 min from stroke onset was reported in the FAST-MAG trial in 2015^53^. More recently, the RIGHT II and MR ASAP trials, conducted in Europe achieved median administration times of 73 min and 63 min with transdermal NG, respectively^41,43^.

In conclusion, our results demonstrate that treatment with NG-NPAs significantly increased LMC-mediated penumbral perfusion, decreased infarct volume, and reduced neurological deficits 24 hours after induction of experimental stroke in hypertensive rats, with poor baseline collateral status. Administration of free NG at the dose equivalent to that in NG-NPAs did not show benefit in enhancing collateral blood flow and instead caused significant hypotension, as is also observed clinically in stroke patients who were administered with free NG. We suggest that selective targeting of the collateral vessels using NG-NPAs based on their unique shear stress profile during stroke may realise the long-recognised potential of using widely available vasodilators to improve stroke outcomes. This would be accomplished via shear targeted delivery to LMCs thereby enhancing collateral flow and penumbral perfusion, while avoiding counteracting effects on blood flow due to systemic vasodilation and hypotension. Ultimately, if these effects are confirmed in patients, there would be potential benefits to “buy time” prior to administration of reperfusion therapies, in addition to providing hope for those ineligible or unsuitable for such therapies.

## MATERIALS AND METHODS

### Animal studies

Spontaneously hypertensive rats (SHR male, 12-14 weeks old n = 56, Wistars, male, 12-14 weeks, n = 6) were used in this study. Experiments conformed to the Animal (Scientific Procedures) Act 1986 (United Kingdom), the National Institutes of Health guidelines for care and use of laboratory animals and were approved by the University of Oxford Animal Ethics Committee, the Home Office (United Kingdom). Experiments were approved by Animal Care and Ethics Committee of the University of Newcastle, Australia (Protocol # A-2020-003 and Protocol #A-2011-131, respectively), in accordance with the requirements of the Australian Code of Practice for the Care and Use of Animals for Scientific Purposes. The studies were conducted, and the article prepared in accordance with the STAIR recommendations^54^ and reported in accordance with the ARRIVE guidelines ^55^. Detailed animal and experimental descriptions are available in the Supplementary Materials.

### Quantification of fluid shear stress in LMCs

Fluid shear stress was measured in the LMCs of normotensive Wistar rats and SHRs by intravenously infusing fluorescent microspheres (1μm in diameter, 0.2% w/v; Molecular Probes) at the rate of 4mL/h and measuring their movement through the LMC between the ACA and MCA through a cranial window using an Olympus BX60 microscope at 10X magnification. LMC flow was recorded using a fluorescent microscope-mounted high-speed camera (Ace U acA720-520um, Basler, Germany or Genie HM640, Teledyne Dalsa, Canada) at 300 frames/s^22,56^. This allowed us to measure LMC blood flow velocity and LMC vessel diameter. Collateral velocity and diameter data have previously been reported for the Wistar and SHR cohort of animals,^22,56^ however, shear stress was not previously analysed. We used these parameters to quantify total shear stress in these collateral vessels using the Equation: τ = γ × η, where τ is shear stress, γ is shear rate and η is viscosity. Shear rate was calculated as 8 × (blood flow velocity) / (vessel diameter). We used published figures for blood viscosity (3 cP).^23^

### Computational fluid dynamic analysis

A 3D cerebrovascular model was generated from computed tomography angiogram (CTA) images of a patient who had suffered an ischaemic stroke, selected from the International Stroke Perfusion Imaging Registry (INSPIRE). These CTA images were segmented using a modelling and segmentation software 3D Slicer (http://www.slicer.org), according to the method adapted from Bateman et al^57^. To reduce computational time, the cerebrovascular system consists of a complete arterial system of one hemisphere and a removed arterial system on the contralateral side (**Supplementary Fig. 1A).** The length of the left and right internal carotid arteries (ICAs) of the model, which were the inlets to the system, were extended to allow the flow at the inlets to be fully developed before entering the cerebrovascular system. To implement an occlusion, a section of the proximal middle cerebral artery was removed (**Supplementary Fig. 1B**). Tetrahedron mesh was used to model the cerebral vasculature due to the complexity of the geometry. Mesh sensitivity analysis was conducted to ensure the convergence of solutions.

Flow conditions were applied to the inlet and outlet vessels according to flow rate values found in Padmos et al^58^. For inlet velocity, the average flow rate passing through the ICA is approximately 260 ml/min. With a measured ICA diameter of 5.156 mm, this amounts to an average inlet velocity of 0.2075 m/s, which was applied to the two inlets of ICAs indicated by blue arrows in (**Supplementary Fig. 1A**). Outlets of the model were grouped into large vessel outlets and collateral outlets. The flow rates at the outlet of large vessels were obtained (**Supplementary** Error! Reference source not found.) and were setup using the target mass flow rate function within ANSYS Fluent (ANSYS, Inc, Canonsburg USA). A gauge pressure of 40 mmHg was set at the outlets of remaining smaller vessels for the occluded model. For the non-occluded model, the outlet pressure at the vessels was set to 0 mmHg. For the wall boundary condition, a rigid arterial wall with a no-slip condition was applied. The shear stress transport fluid model was applied to simulate the complex cerebrovascular model with large variations in geometry.

Two LMC vessels were extracted from the full cerebrovascular model using CREO (PTC, Boston USA). The two collateral vessels were meshed with 429,425 and 623,531 elements respectively to analyse their wall shear stress with higher details and accuracy. The average inlet velocity to each collateral was obtained from the simulated results of the full cerebrovascular models. The boundary conditions and average wall shear stress for each collateral vessel can be found in **Supplementary** Error! Reference source not found..

### NG-NPA preparation and characterisation

NG-NPAs were fabricated in a two-step process. First, we encapsulated NG (USP grade, American Regent) into polylactic-glycolic acid (PLGA; 50:50) nanoparticles via single emulsion technique using a probe sonicator (Q-Sonica modelQ700 with a probe model CL-334). This emulsion was purified by dialysis (Biotech CE Dialysis Tubing; MWCO 100kD) overnight. Dynamic Light Scattering (DLS) was used to determine the size of these NG loaded nanoparticles (Malvern instruments, UK).

The second step was the production of NG-NPAs via spray drying. NG loaded nanoparticles synthesized in the first step were brought to 5 mg/ml concentration by adding Milli-Q water and then mixed with 1 mg/ml L-leucine (Spectrum Chemicals & Laboratory Products, CA). This aqueous leucine nanoparticle suspension was mixed with ethanol at a ratio of 1:1.5 and immediately spray dried using a Buchi 290 spray dryer (Büchi Labortechnik AG, Flawil, Switzerland) maintaining the inlet temperature between 105-110°C, outlet temperature below 40°C, and the feed rate 5 mL/min.

The size of the NG-NPAs were determined by using laser diffraction (Mastersizer, Malvern Instruments, UK). Ultrasound sonication assay was used to quantify shear disaggregation of the NG-NPAs as previously described^59^ and NG loading in the NG-NPAs was measured by using LC-MS.

### Surgical Procedures

#### Laser Doppler Flowmetry (LDF)

LDF was used to measure changes in cerebral blood flow (CBF) in the MCA territory. The LDF probe (Moor Instruments, UK) was inserted into the hollow PEEK screw (Solid Spot LLC, Santa Clara, CA, USA) located +5 mm lateral of midline and -2 mm posterior to Bregma in the right parietal bone.

#### Laser Speckle Contrast Imaging (LSCI)

LSCI (RWD Life Sciences, China) through bilateral thinned-skull cranial windows was used to measure changes in CBF in the distal middle cerebral arterial territory supplied by retrograde collateral flow on the right (stroke) side of the brain and in the matching region contralaterally. Bilateral cranial windows (5 mm x 5 mm) were created extending posterolaterally from Bregma 1mm posterior, 1mm lateral, according to our published method^22^. Regions of interest (ROI) on laser speckle imaging software were placed 2 mm posterior and 3 mm lateral from Bregma to measure changes in CBF within the the terminal MCA perfusion territory, supplied by LMC (**Fig. 1a**). Placement of ROI are based off cerebrovascular casting studies^60,61^ and have been validated using magnetic resonance imaging^62^ and hydrogen clearance^63^. A matching ROI was placed in the homotypic contralateral region. To measure vein diameters, LSCI with higher magnification was used to zoom in on the right (stroke) side of the brain. The laser speckle contrast imaging data was collected at pre-stroke baseline, post-stroke pre-infusion, and then continuously during 40 min drug infusion. Perfusion data analysis and vein diameter analysis was performed by an investigator blind to treatment groups, using LSCI analysis software (RWD Life Sciences, China).

#### Intracranial Pressure (ICP)

An ICP probe (OpSens Fiber Optic Pressure Sensors, Canada) was inserted epidurally through a hollow poly-ethyl ether ketone (PEEK) screw in the left frontal bone, according to our previously described method^64^. The screw was secured using ethyl 2-cyanacrylate (Super Glue Ultra-Fast liquid, UHU, Australia) and biocompatible caulking material (Silagum, Gunz Dental, Germany).

#### Middle cerebral artery occlusion (MCAo)

MCAo was induced according to our established protocol^65^. To summarize, a silicone-tipped monofilament was inserted into the external carotid artery and advanced 20 mm up the internal carotid artery until resistance was felt and a drop in perfusion (>70%) on LDF was observed, indicating occlusion of the origin of the middle cerebral artery. The filament was retracted after 70 min to produce recanalization.

### Study design

*Study I -* In Study I we investigated the shear stress in LMC during experimental stroke in Wistar rats and SHRs. Detailed methods and collateral velocity and diameter data have previously been reported for the Wistar rats and SHRs^22,56^. However, the shear stress was not previously analysed. Rats underwent baseline recordings of anterior cerebral artery-middle cerebral artery (ACA–MCA) LMC blood flow velocity and diameter before induction of experimental stroke (90 min MCAo, n = 6 Wistars and n = 5 SHRs). LMC flow velocity and diameter recordings were then taken every 10 min throughout occlusion. Animals underwent baseline recordings of anterior cerebral artery-middle cerebral artery (ACA–MCA) LMC blood flow velocity and diameter before induction of experimental stroke (70 min MCAo, n = 6). Collateral flow velocity and diameter recordings were then taken every 10 min throughout occlusion and an average was taken for the duration of the occlusion. Shear stress was then calculated using blood flow velocity, diameter, and viscosity^23^. For shear stress calculations see Methods in the Supplementary Materials. (A detailed experimental timeline can be found in **Supplementary Fig. 2A**)

### Study II

In study II we determined the effect of NG-NPAs on LMC-mediated penumbral perfusion during stroke in SHRs. Pre-stroke baseline measurements of LMC perfusion, contralateral perfusion and mean arterial pressure (MAP), were taken, and then rats were subjected to 70 min of MCAo. A post-occlusion/pre-drug infusion baseline measurement of LMC and contralateral perfusion, and MAP was conducted 25 min after MCAo. Animals were then randomized by sealed numbered envelope to receive intravenous infusion of either Blank-NPAs (control, 4mg in 2ml of saline, n = 7) or NG-NPAs (50μg nitroglycerin in 4mg NPAs in 2ml of saline at 2.8-3ml/h = 4μg/kg/min, n = 7), for 40 min. Dose and sample sizes were based on results from a preliminary study shown in the Supplementary Materials. All treatments were administered by a surgeon who was blind to treatment allocation at the drug administration stage and remained blinded during all subsequent analysis of results. LMC perfusion, contralateral perfusion and MAP recordings continued until thread withdrawal (recanalization) 70 min post-MCAo. Following 24 h of recovery, animals were tested for stroke-induced neurological deficits and brains were collected for infarct volume, performed using both Triphenyltetrazolium (TTC) staining and haematoxylin and eosin (H&E) histology per our standard technique^66^ (detailed methods in Supplementary Material and experimental timeline in **Supplementary Fig. 2B**).

### Study III

In study III we determined the effect of NG-NPAs on cortical vein diameter and ICP. The experimental protocol was the same as study II, except that higher magnification of laser speckle contrast imaging was used and ICP was also measured (detailed timeline in **Supplementary Fig. 2C**). Laser speckle imaging was performed through the cranial window prior to induction of MCAo and during infusion of NG-NPAs (n = 6) or Blank-NPAs (n = 6). Changes in cortical veins diameters and ICP was monitored at pre-stroke baseline and post-stroke at pre-infusion baseline and throughout 40 min drug infusion. ICP data was recorded for the NG-NPAs (n = 3) and blank-NPAs (n = 4) groups (detailed timeline in **Supplementary Fig. 2C**).

### Study IV

In Study IV we determined the effect of free nitroglycerin (free-NG) on LMC perfusion and MAP during stroke. The protocol was as for Studies II and III, except animals were randomized to receive intravenous infusion of saline (n = 6) or infusion of nitroglycerin (0.25 μg/µl at 300µl/h = 4 μg/kg/min of NG, n = 6 or 0.5 μg/µl at 300µl/h = 40 μg/kg/min of NG, n = 4), commencing 25 min after MCAo (detailed timeline in **Supplementary Fig. 1D**).

### Exclusion Criteria and Statistical Analysis

Subarachnoid haemorrhage, experimental complications and ≤ 70% drop on laser Doppler flowmetry at time of middle cerebral artery occlusion were pre-specified exclusion criteria. D’Agostino and Pearson omnibus normality tests were performed on all data. When comparing two-independent variables, normally distributed data was compared using unpaired student’s t-test and non-normally distributed data was compared using Mann Whitney-U test. When comparing how a response is affected by two factors where one of the factors was repeated; independence, normality, and sphericity was assumed, and variables were compared using Repeated measures two-way ANOVA with Sidak’s and Dunnett’s multiple comparisons test between groups. Correlations were classified as tiny or < 0:05, very small (0:05 < = r < 0:1), small (0:1< = r < 0:2), medium (0:2< = r < 0:3), large (0:3< = r < 0:4), or very large (r > =0:4) according to Funder and Ozer’s criteria^67^. Detailed description of sample size calculations for LMC perfusion are available in the Supplementary Materials. Statistical tests used for each analysis are reported in figure legends. For ANOVAs, overall ANOVA results are reported in figure legends and results of post hoc tests are reported in the results text and are represented in each figure. Significant differences were accepted at the level of p < 0.05; data are presented as mean ± SEM.

## Acknowledgements

This work was funded by Oxford University Medical Sciences Internal Fund Pump Priming award (DB, DEI, AMB), National Health and Medical Research Council of Australia APP1182153 (DB, DEI, NJS), and the Wyss Institute for Biologically Inspired Engineering at Harvard University (DEI).

## Author Contributions

Conceptualization: AMB, NJS, DEI, DJB; Methodology: ML, SA, RJH, KM, DP, DO, YKY, AC, NH, SJ, PGB, FM, CGE, DB, OU, NK, DJB, DEI; Investigation: ML, NJS, DJB; Visualization: ML, NJS, DJB; Funding acquisition: YC, AMB, NJS, DEI, DJB; Project administration: DJB; Supervision: NJS, DJB, DEI; Writing – original draft: ML; Writing – review & editing: ML, SA, RJH, KM, DP, DO, OU, DB, NK, YC, AMB, NJS, DJB, DEI.

## Data and materials availability

The data that support the findings of this study are available from the corresponding author upon reasonable request.

