## Supplementary information for "Shear stress targeted delivery of nitroglycerin to brain collaterals improves ischaemic stroke outcome"

### SUPPLEMENTARY METHODS

#### Pilot Study

##### *Animals*

Animal experiments were performed on male Spontaneously Hypertensive Rats (SHRs) weighing 300-350 g (Harlan, Bicester, UK). All animals were housed in a 12 h light/dark cycle and had ad libitum access to food and water prior to experiments. All procedures conformed to the Animal (Scientific Procedures) Act 1986 (UK) and the National Institutes of Health guidelines for care and use of laboratory animals and were approved by the University of Oxford Animal Ethics Committee, the Home Office (UK).

##### *Anaesthesia and Monitoring*

Rats were anesthetized with 5% isoflurane in $O_2/N_2$ (1:3) and maintained with 1% to 2% isoflurane. Core temperature was maintained at 37˚C by a thermocouple rectal probe and warming plate (Harvard Apparatus, UK). Incision sites were shaved, cleaned, and injected subcutaneously with 2 mg/kg 0.05% Bupivacaine (Aspen, UK). A femoral arterial line was used for continuous arterial pressure monitoring.

***Experimental Stroke Model***

Rats underwent middle cerebral artery occlusion (MCAo/stroke) using the silicone-tipped intraluminal thread occlusion method (as set out in Spratt *et al*.[1]), using 4-0 monofilament occluding threads with 4 mm length x 0.35 mm diameter silicone tips. The filament was advanced into the right external carotid artery stump and up the internal carotid artery to occlude the origin of the right MCA.

***Measurements of changes in core and collateral blood flow using multi-site dual laser Doppler probes***

Laser Doppler flowmetry (LDF) (Oxford Optronix, Oxford, UK) was used to measure changes in cerebral blood flow (CBF) in both the MCA and collateral arterial territories using dual probes. The animal's head was secured with ear bars in a stereotaxic frame. Probe 1 was placed +4 mm lateral of midline and -2 mm posterior of Bregma to measure changes in core MCA CBF. Probe 2 was placed +3 mm lateral of midline and +2 mm anterior of Bregma to measure changes in CBF supplied by collateral vessels within the border zone between the anterior cerebral artery (ACA) and MCA perfusion territories, as previously described in Beard et al[2]. MCAo was confirmed by >70% decrease in LDF signal from baseline in Probe 1.

***Drug administration***

Prior to MCAo, the femoral vein was cannulated with 2-French silicon tubing. Animals were randomized to receive intravenous bolus and then infusion of blank nanoparticle aggregates (Blank-NPA, 1 mg in Saline n=6) or bolus and then infusion of nitroglycerin nanoparticle aggregates (NG-NPAs, 12.5 μg nitroglycerin in 1 mg NPA in saline, n= 6) I.V, 30 minutes after MCAo. Collateral perfusion was measured as a % change from pre-injection baseline.

### **Main Study**

#### ***Animals***

All animal experiments were performed on male SHRs (12-14 weeks old, weighing 270-320g; ARC breeding facility in Perth, Australia). All animals were housed in a 12 h light/dark cycle and had ad libitum access to food and water prior to experiments. All procedures were approved bythe Animal Care and Ethics Committee of the University of Newcastle (Protocols # A-2020-003) and in accordance with the requirements of the Australian Code of Practice for the Care and Use of Animals for Scientific Purposes. The relevant Stroke Treatment Academic Industry Roundtable (STAIR) recommendations for preclinical research (randomization, blinded assessment, physiological monitoring) were strictly followed[3]. The manuscript was prepared in accordance with the ARRIVE guidelines[4].

#### ***Anaesthesia and monitoring***

In all studies, rats were anesthetized with 5% isoflurane using an induction chamber in 50:50 $O_2/N_2$ and maintained with 1% to 2% isoflurane. Incision sites were shaved, disinfected, and injected with local anaesthetic (2mg/kg 0.05% Bupivacaine, s.c.).

During surgery and imaging, temperature was maintained at 37° by a thermocouple rectal probe and warming plate (RET-2, Physitemp Instruments Inc, USA). Systemic blood pressure and heart rate was continuously monitored via catheter placed in the right femoral artery. Blood samples (0.1mL) were collected before and after drug infusion for blood gases analysis (i-STAT, Abbot, USA). In study II, after stroke surgery, animals were injected subcutaneously with saline (2x 2.5ml) to prevent dehydration and placed in their cages with free access to soft sweetened food and water and recovered for 24 hours.

***Collateral Shear Stress Calculation***

In study I, Collateral velocity and diameter data have previously been reported for the Wistar and SHR cohort of animals[5,6], however, shear stress was not previously analysed. In brief, fluorescently labelled microspheres (1 μm diameter, 0.2% w/v) (Molecular Probes) were continuously infused through the jugular line (4mL/h) and could be seen transiting collateral vessels on the pial surface through a closed cranial window. The velocity of the microspheres and collateral vessel diameter were recorded at 300 frames per second with a digital camera (Ace U acA720-520um, Basler, Germany or Genie HM640, Teledyne Dalsa, Canada) connected to a x10 objective fluorescent microscope (BX60, Olympus, Japan). Shear stress was calculated as $\tau = \gamma \times \eta$, where $\tau$ is shear stress, $\gamma$ is shear rate and $\eta$ is viscosity. Shear rate was calculated as 8 × (microsphere velocity) / (vessel diameter). Viscosity was set to 3 cP[7].

***Histological examination***

In study II, animals were euthanized 24 h post-MCAo and perfused transcardially with saline. Brains were removed and sliced to 2mm coronal section using rat brain matrix. Triphenyltetrazolium (TTC) (Sigma Aldrich, Missouri, USA) was used to stain for infarct volume determination. The brain slices were incubated for 12 minutes at 37°C in 2% TTC. TTC was used for early confirmation of infarct and initial infarct analysis. Infarct was identified as a white area of each section and measured with ImageJ software (National Institutes of Health, Bethesda, MD)[8]. Haematoxylin and eosin (H&E) staining was then used on the same tissue for further confirmation of infarct and infarct volume quantification. The brains were cut into 10-micron coronal sections and stained with haemotoxylin–eosin. Sections were imaged at x40 using an Aperio slide scanner (Aperio Technologies Inc.USA) and three regions of interest (ROIs) were traced using Aperio's imagescope software: the stroke hemisphere (ipsilateral to MCA occlusion), the non-stroke hemisphere (contralateral hemisphere ROI) and the infarct area (infarct ROI). Tracing the infarct area under high-power magnification, instead of low power, prevented the under- or over-estimation of the infarct area. Microscopically, infarct areas were represented by complete neuronal eosinophilia, with condensed nuclei, cytoplasmic scalloping and no visible nucleolus. Areas of selective neuronal necroses were not included in the infarct ROI. Macroscopically, the infarct area was represented by the area of pallor. The infarct volumes for each animal were calculated by averaging the infarct area between adjacent slices and multiplying by the distance between slices. Volumes were then corrected for edema using the formula: corrected infarct volume ($mm^3$) = infarct volume x (contralateral volume/ipsilateral volume).

***Neurological Deficit Testing***

In study II at 24 hours post-stroke, the forelimb flexion, torso twist and lateral push tests were used to determine neurological deficit score, a total neurologic deficit score given out of 6 (higher score indicating greater deficit)[9].

***Detailed Sample Size Calculations***

The primary outcome measure was changes in LMC perfusion in Study II. All other outcomes are secondary outcomes and considered exploratory. In our preliminary study conducted in SHR rats we found that NG-NPA treatment increased collateral perfusion by 42 ± 27 % vs. -3 ± 7% for the NG-NPA group (**Supplementary Fig. 3**). These values give a Cohen's d effect size of 2.25. Therefore, we required 7 animals per treatment group to be able to reject the null hypothesis that NG-NPAs do not enhance collateral flow with probability (power) 0.95. The type I error probability associated with the test of this null hypothesis (alpha) was 0.05.

### <u>SUPPLEMENTARY RESULTS</u>

#### NG-NPA Collateral Pilot Study

NG-NPAs significantly increased LMC perfusion (collateral blood flow). LMC perfusion was increased relative to Blank-NPAs and pre-infusion baseline between 5- and 10-minutes post-infusion (5 minutes, NG-NPA: 42.4 ± 19 vs. Blank-NPAs: -3.9 ± 3.1 % of pre-infusion baseline, $p < 0.0001$, **Supplementary Fig. 3A**). NG-NPAs significantly increased the average LMC perfusion during infusion (NG-NPA:

20.4 ± 3.3 vs. Blank-NPAs: -2.4 ± 2.2 % pre-infusion baseline, $p = 0.0002$, **Supplementary Fig. 3B**).

**Study II Exclusions and Physiological Variables**

A total of 9 animals were excluded. Reasons for exclusions were experimental complications (n = 6), lack of sufficient LDF drop (n=1), and subarachnoid haemorrhage detected on laser speckle imaging and confirmed at post-mortem (n=2). An additional NG-NPA animal was excluded from infarct volume analysis due to an experimental complication just after successful reperfusion and lack of subarachnoid haemorrhage, so it was decided that the LMC perfusion and mean arterial pressure data from this animal (collected before reperfusion) could be included for analysis. Therefore, a total of 14 animals (NG-NPA n = 7, Blank-NPA n =7) were included in the final LMC perfusion analyses and 13 animals (NG-NPA n = 6, Blank-NPA n =7) were included in infarct analysis. Physiological variables (mean arterial blood pressure, $pO_2$, $pCO_2$) were stable throughout the experimental protocol with no significant differences in any variable between NG-NPAs and Blank-NPAs groups (**Table 1**).

**Study II TTC Infarct Analysis**

NG-NPA significantly reduced TTC-determined infarct volume at 24 h following stroke (NG-NPA: 75 ± 9.7 vs. Blank-NPAs: 126.5 ± 13.6 $mm^3$, $p = 0.01$, **Supplementary Fig. 4A**). There was a very large, significant inverse correlation between average LMC perfusion and infarct volume ($r = -0.7$, $p = 0.007$, **Supplementary Fig. 4B**).

### SUPPLEMENTARY FIGURES

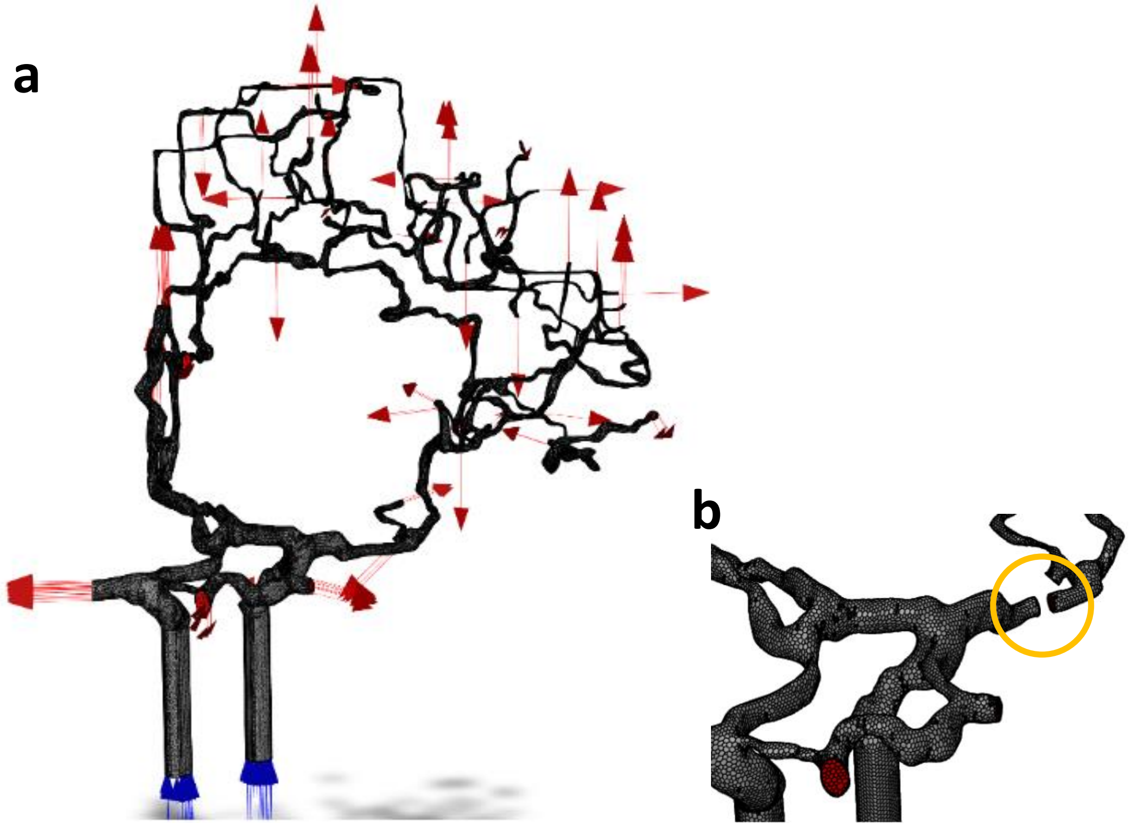


**Supplementary Figure 1**. **(a)** The cerebrovascular system with an arterial system of one hemisphere. Blue arrows indicate inlets and red arrows indicate outlets. **(b)** Removal of a section in MCA to simulate occlusion.

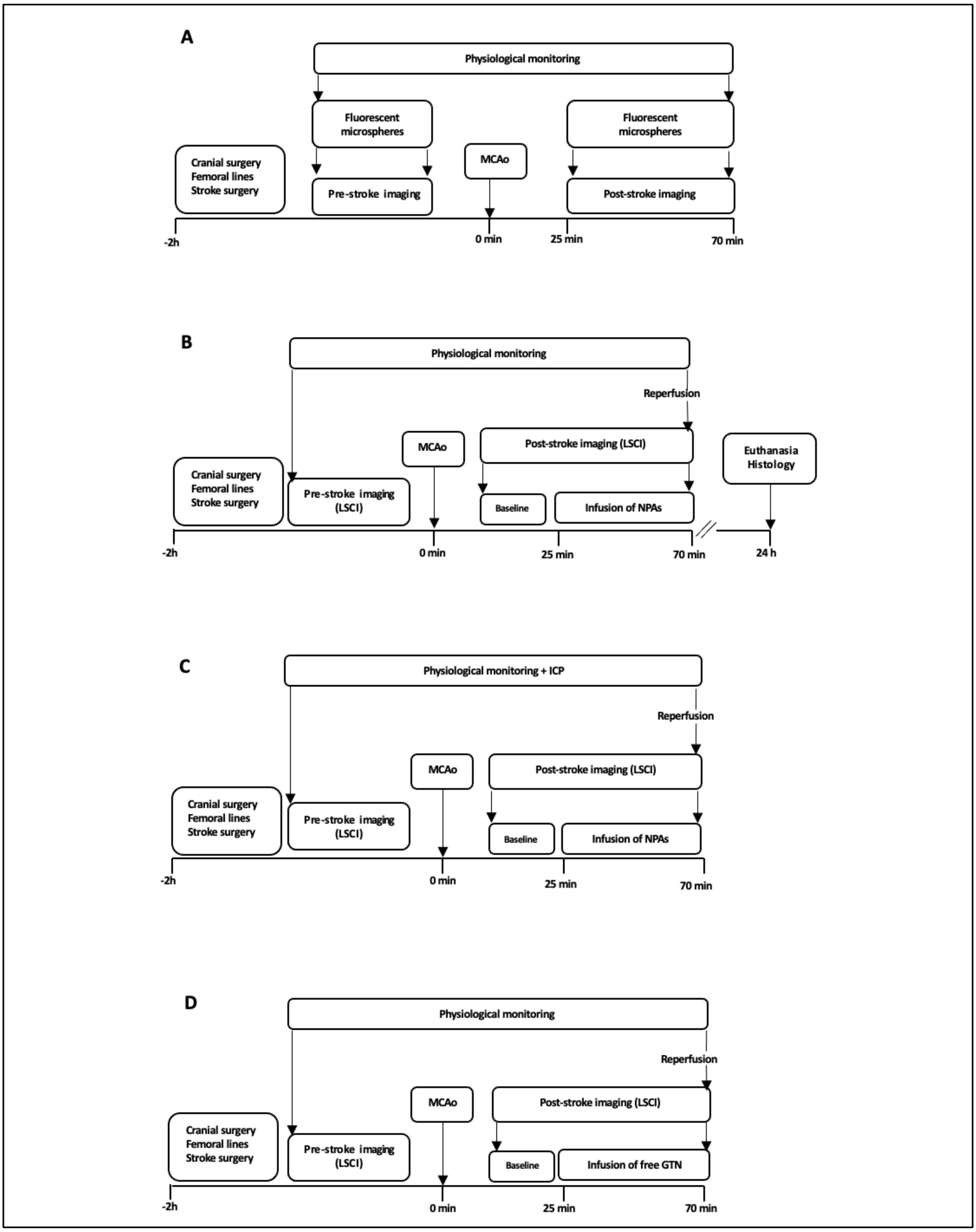


**Supplementary Figure 2.** Experimental protocols. Physiological parameters were monitored throughout the procedure for all experiments. Baseline perfusion imaging recordings were taken for 5 minutes prior to starting MCAo for all experiments. 0h indicates the moment of vessel occlusion. Fluorescent microspheres were infused during baseline imaging and 25 minutes after MCAo for 40 minutes. Recordings were taken every 10 minutes throughout the occlusion **(a)**. Treatment with NPAs was initiated 25min post occlusion until reperfusion at 70 min. Infarct volume was assessed at 24 hours **(b)** ICP was measured before MCAo, pre-infusion and during 40 minutes of drug infusion **(c).** Treatment with free-NG was initiated 25min post-MCAo until reperfusion at 70min **(d).** Recordings on LSCI were taken every 2min for 40 min during NG-NPA/free NG infusion. MCAo, middle cerebral artery occlusion; LSCI, Laser Speckle Contrast Imaging; ICP, intracranial pressure; NPAs, nanoparticles; Free NG, free nitroglycerin.

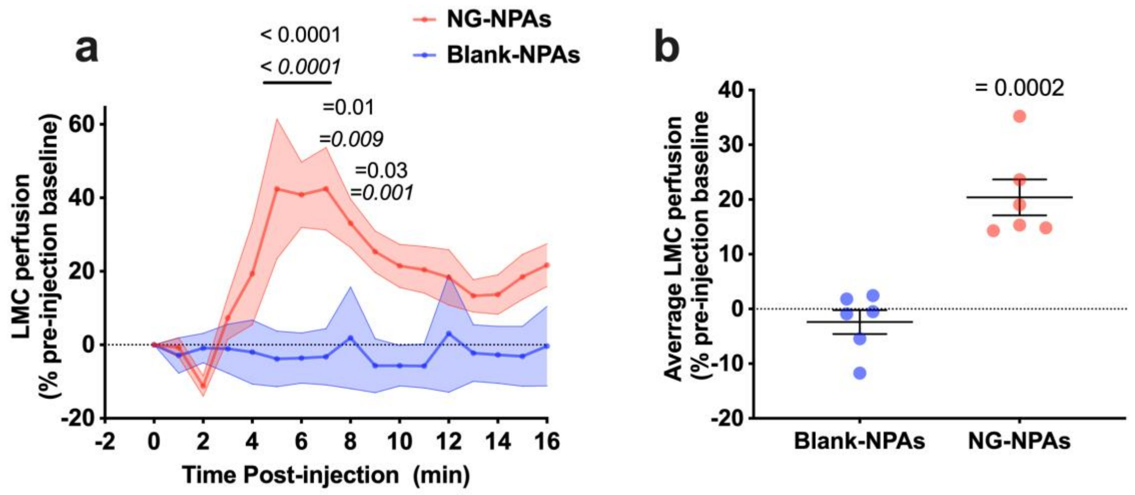


**Supplementary Figure 3. (a)** LMC perfusion in the stroke region in NG-NPA (red) treated vs. Blank-NPA (Blue) treated SHRs, calculated as a % change from pre-drug injection baseline. We conducted a repeated measure 2-Way ANOVA to assess the effect of NG-NPA vs. Blank-NPA treatment over time. $F (1, 10) = 33.1$, $p = 0.0002$ for treatment, $F (16, 160) = 3.2$, $p < 0.0001$ for time, $F (16, 160) = 3.7$, $p < 0.0001$ for interaction. Sidak's post-test was used to compare NG-NPA vs. Blank-NPAs at different time-points (p values on graph in italics, $p < 0.0001$ 5-7 minutes, $p < 0.001$ 8-9 minutes and $p < 0.05$ at 10 minues) . Dunnett's post-test was used to compare each time-point back to pre-infusion baseline in NG-NPA and Blank NPA groups (NG-NPA: $p<0.05$ at 8-9-minutes; $p< 0.0001$ between  5 and 7 minutes).

(**a**) LMC perfusion following NG-NPA vs Blank-NPA in SHR in pilot experiment. **(b)** Average LMC perfusion in the stroke region. We used an un-paired t-test to assess differences between NG-NPAs (red dots) vs. Blank-NPAs (blue dots). T (10) = 5.8, p = 0.0002 vs. Blank-NPAs. Values represent mean ± SEM.

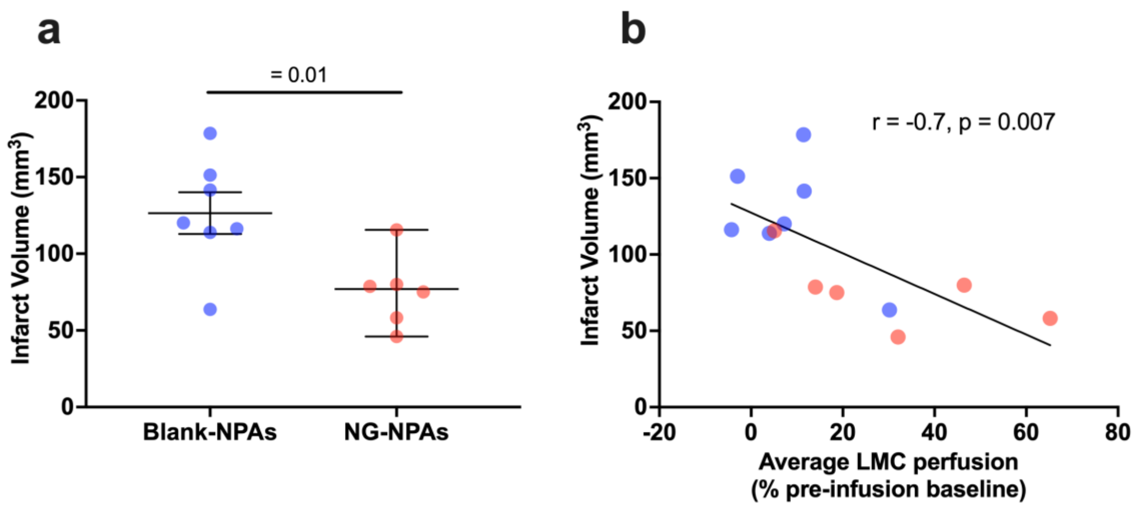


**Supplementary Figure 4.** (**a**) Infarct volume was assessed using TTC at 24 hours after stroke onset. We used an un-paired t-test to differences between NG-NPAs (red dots) vs. Blank-NPAs (blue dots). T (11) = 2.9, p = 0.01 vs. Blank-NPAs. **(b)** Pearson's correlation between infarct volume and average LMC perfusion.

**Supplementary Tables**

***Supplementary Table 1**. Large vessel flow rates, measured diameters and mass flow rates.*

| Large Vessel | Volumetric Flow Rate [ml/min] | Measured Diameter [mm] | Inlet velocity [m/s] | Outlet mass flow rate [kg/s] |
|---|---|---|---|---|
| ICA | 260 | 5.156 | 0.2075 | |
| MCA | 146 | 2.981 | | 0.0026 |
| PCA | 54 | 3.08 | | 0.000954 |
| ACA | 82 | 1.908 | | 0.0014 |

**Supplementary Table 2.** Boundary conditions and average wall shear stress of collateral vessels.

| | Inlet velocity [m/s] | | Wall Shear Stress [dynes/cm$^2$] | |
|---|---|---|---|---|
| | Non-occluded | Occluded | Non-occluded | Occluded |
| Collateral 1 | 0.041 | 0.89 | 3.94 | 307.61 |
| Collateral 2 | 0.04 | 0.91 | 15.93 | 348.65 |
